# Transcription Factor Gene *Pea3* Regulates Erectile Function in Mice

**DOI:** 10.1101/2021.11.15.468751

**Authors:** Jarret A.P. Weinrich, Aanchal Tyagi, Megan C. Kenney, Richard J. DiCasoli, Julia A. Kaltschmidt

**Author notes:** Department of Anatomy, University of California San Francisco, San Francisco, CA 94158, USA. AUTHOR CONTRIBUTIONS JAPW performed sexual behavior assays and wrote MATLAB analysis code. AT and MCK quantified animal behaviors. RJD provided technical support. JAPW and JAK designed the study, interpreted results and wrote the manuscript.

## Abstract

**Background:** Male mice with homozygous loss of function mutations of the *ETS* transcription factor gene *Pea3* (*Pea3* null) are infertile due to their inability to deposit semen plugs, however the specific deficits in male sexual behaviors that drive this phenotype are unknown.

**Aim:** To investigate the regulatory role of the *Pea3* gene in organizing gross sexual behaviors and erectile functioning during active copulation.

**Methods:** The copulatory behavior of male mice (*Pea3* null and control) with hormonally primed ovariectomized females was monitored via high-speed and high-resolution digital videography to assess for differences in female-directed social behaviors, gross sexual behaviors (mounting, thrusting), and erectile and ejaculatory function.

**Outcomes:** *Pea3* null male mice have dramatically reduced erectile function during sexual intercourse, however other aspects of male sexual behaviors are largely intact.

**Results:** *Pea3* null male mice exhibit greatly reduced erectile function, with 44% of males displaying no visible erections during mounting behaviors, and none achieving sustained erections. As such, *Pea3* null males are incapable of intromission, and semen plug deposition, despite displaying largely normal female-directed social behaviors, mounting behaviors, and ejaculatory grasping behavior. Additionally, the coordination of the timing of thrusting trains is impaired in *Pea3* null males.

**Clinical Implications:** The identification of the transcription factor *Pea3* in regulating erectile function in mice may provide a useful target for understanding the genetics of male sexual dysfunction in human patients.

**Strengths and Limitations:** High-speed and high-resolution videography allows for a detailed analysis of male sexual behaviors and erectile functioning in *Pea3* null and control mice. How disruption of the *Pea3* gene translates to erectile dysfunction is still unknown.

**Conclusion:** The transcription factor gene *Pea3* regulates the ability to achieve and maintain erections in male mice.

## INTRODUCTION

For males, successful copulation involves the tight coordination of context-dependent social behaviors, erectile functioning, and ejaculation. Approximately 20-50% of men may suffer sexual dysfunction^1,2^, which can range from diminished sexual arousal and erectile function to the complete absence of erections and ejaculations^3^. Human genetic factors for erectile dysfunction have recently been discovered^4,5^, and several genetic mouse models of male sexual dysfunction have been identified that alter the propensity to engage in sexual behaviors, diminish erectile function, or abolish gross ejaculatory behaviors. These mouse models have been linked to changes in the central nervous system^6–9^, breakdown of structural integrity of the penile epithelia^10,11^, disruption of cellular signaling between the autonomic nervous system and penile vasculature and epithelia^12–14^, and diminished contractile ability of the vas deferens^15^. However, many of these mouse models are at least minimally fertile, and thus mutant males possess the ability to successfully copulate. Therefore, the molecular underpinnings of absent erectile or ejaculatory function are still not well defined.

Male mice with a homozygous loss of function mutation for the *ETS* transcription factor gene *Pea3* (*Pea3* null) are completely infertile due to an inability to deposit semen plugs during copulation^16^. In adult mice, *Pea3* is expressed within male reproductive organs (testis and epididymis), suggesting a possible role for *Pea3* in spermatogenesis or maturation^16,17^. However, *Pea3* null males produce sperm capable of fertilization using *in vitro* assays^16^. Similarly, *Pea3* null mice do not exhibit any identifiable histological deficits in penile tissue, no loss of corpus cavernosum contractile ability in *ex vivo* assays, and exhibit an intact capability for reflex erections, suggesting that erectile functioning is intact^16^. Furthermore, during the initial characterization of the *Pea3* null sexual behavior phenotype, Laing *et al* witnessed *Pea3* null males performing mounting behaviors at the start of overnight mating assays, suggesting that both appetitive and consummatory male sexual behaviors are at least partially intact^16^. A systematic characterization of male sexual behaviors, including direct observation of the penis during copulation, of *Pea3* null male mice has yet to be performed. The specific behavioral deficits that accompany, and potentially drive, the abolishment of plug deposition in male mice with loss of *Pea3* gene are unknown.

Here, we hypothesized that the *Pea3* null male infertility is due to (1) severe erectile dysfunction and/or (2) the complete loss of ejaculatory function. To ascertain deficiencies in *Pea3* null male-specific female-directed social behaviors, gross sexual behaviors, and erectile and ejaculatory functioning, we employed a sexual behavior arena^18^ that allowed for direct observation of erectile function and monitored mouse behaviors using high speed and high resolution digital video recording, which was followed by a detailed post hoc analysis. Surprisingly, we found that loss of *Pea3* leads to greatly diminished erectile functioning, with many mice displaying no visible erectile activity, and those that do only produce brief, rare, and poorly timed periods of penile tumescence. Additionally, we found that the coordination of hip thrusting during sexual behavior is perturbed in *Pea3* null males. These results indicate that the transcription factor *Pea3* is an important regulator of erectile functioning during copulation in mice.

## MATERIALS AND METHODS

### Animal husbandry

All mouse husbandry and surgical procedures strictly adhered to the regulatory standards of the Institutional Animal Care and Use Committee of Memorial Sloan Kettering Cancer Center (MSKCC; protocol 08-06-009). The following mouse strains were used in this study: Pea3^NLZ^ mice (also known as Etv4^tm1Arbr^)^19^ and ovariectomized c57BL6/J females (Stock: 00064; Jackson Laboratories). Ovariectomized females were procured from Jackson Laboratories.

### Preparing for behavioral testing

Sexual behavior was conducted as in Park (2011)^18^ and Juntti *et al* (2010)^7^. Briefly, male sexual behavior was assessed in a closed, rectangular plexiglass arena with a clear plexiglass bottom, below which an angled mirror was placed to allow for viewing of the mouse underside during the behavioral assay^18^. Both male and female mice were prepared for behavioral testing days in advance. Males were singly housed for at least one week before testing, and contact with cages (i.e., cleaning, etc.) were minimized. On each of the two days before testing, ovariectomized females were hormonally primed with a subcutaneous injection of 1 microgram of estradiol benzoate in sesame oil. 4 to 6 hours before testing, ovariectomized females were hormonally primed with a subcutaneous injection of 100 micrograms of progesterone in sesame oil. One hour before recording, males were transferred to the plexiglass arena and allowed to acclimate to the testing environment. Testing was only performed during the dark phase of the light/dark cycle set by MSKCC animal facility staff.

### Behavioral testing

Following acclimation to the testing arena, digital recording was initiated and the hormonally primed female was introduced to the testing arena. Behavior was recorded with a Photonfocus 2048×1088C Series camera with NORPIX Streampix software, at a frame capture rate of 60 frames per second. If males did not commence sexual behavior within 15 minutes of introducing the female, the female was exchanged for an untested female until the male was introduced to 3 females. Behavior was recorded until the male ejaculated or within one hour of introducing the female to the cage, whichever occurred first. Following the termination of testing, the females were checked for semen plugs and then placed into a separate cage from non-tested females. Males were placed back into their home cage. The behavioral arena was carefully washed with soap and water and dried before the next use.

### Video data quantification

Behavioral videos were manually scored within NORPIX Streampix software, with the scorers blind to the genotype of the male in each recording. Relevant data were recorded in excel and when possible the frame number for the start and end of the specific behaviors analyzed were included. Sexual behaviors were classified as follows: (a) mounting: the male positions himself parallel to the female’s body axis and grasps the female’s flank with his forearms, (b) thrusting: repetitive oscillations of the male’s hips while in the mounting position, (c) intromission: insertion of the penis into the vagina, and (d) ejaculatory grasping: during to the male orgasm, the male tightly grasps the female and falls over onto one side. The following data were recorded: introduction of the female to the behavioral arena (time), start of each mount (time), the presence or absence of visible erections, the number of intromissions per mount, and the start and end of ejaculatory grasping (time). For the first and last mount of each recording, the thrust timing was recorded as the frame number from the beginning of one thrust to the beginning of the next thrust.

Social behaviors were assessed as in Fairless *et al* (2013)^20^. Briefly, the interaction of the male with the female was assessed by measuring the cumulative time the male spent sniffing the nose, body, or anogenital regions of the female. The time the male spent grooming the female was also measured. The start and end time of each behavior was recorded to determine the cumulative time for each behavior.

### Data analysis and statistics

Data were processed and analyzed in Mathwork’s MATLAB (R2017a) software. Intromission-like thrusts were defined as any thrust that took between 0.2 to 1 seconds to complete. To determine the number of intromission-like thrusts, the number of thrusts that occurred within this window were counted. Probability distributions of thrust timings were generated using the KSdensity function within MATLAB. Where indicated, data were processed by Fisher’s exact test, student’s t-test or two-way ANOVA performed with either MATLAB or Prism statistical analysis software. The threshold for significance for all statistical tests were set at p < 0.05, and indicators of significance levels were as follows: ns (no significance) when p > 0.05; (*) when p < 0.05; (**) when p < 0.01; (***) when p < 0.001; and (****) when p < 0.0001. Where noted, data in figure legends are reported as the mean value ± the standard error of the mean.

## RESULTS

### FEMALE-DIRECTED SOCIAL BEHAVIORS UNAFFECTED IN PEA3 NULL MALES

First, we set out to determine whether the inability of *Pea3* null mice to set semen plugs was due to a decreased or abolished drive to interact with receptive females, thereby reducing non-sexual aspects of mating behavior. To this end, the duration of time males spent sniffing various regions of the female, specifically the nose, body, and anogenital region, and additionally, the duration of time of female-directed grooming, was quantified^20^. These behaviors were measured for 15 minutes or until the first mount occurred. When compared to control mice, we found that the cumulative time spent performing these behaviors was unaffected in *Pea3* null mice, both when measured by total time (Figure 1A) and percentage of time (Figure 1B). Therefore, decreased female-directed behaviors do not underlie the loss of the ability of *Pea3* null males to deposit semen plugs.

**Figure 1.**
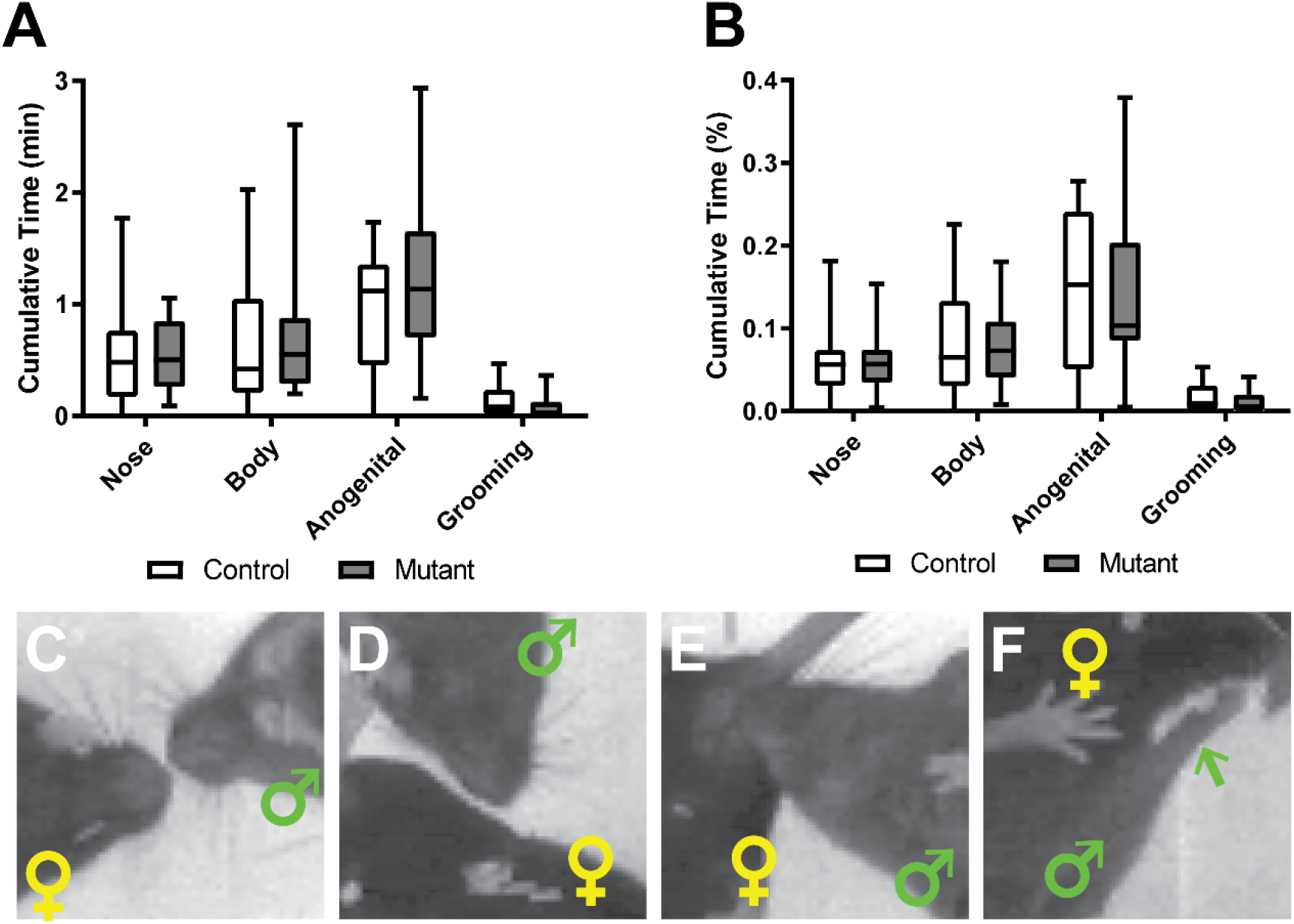
Female-directed social behaviors are unaffected in *Pea3* null mice. (A) The cumulative time spent performing social behaviors in unaffected in *Pea3* null vs control males (Two-way ANOVA, p_genotype_ < 0.5876). (B) The overall percentage of time spent performing social behaviors in unaffected in *Pea3* null vs control males (Two-way ANOVA, p_genotype_ < 0.8983). Representative images of active social behaviors directed to the female nose (C), body (D), and anogenital (E), and grooming (F) behaviors. Male and female positions are marked by green and yellow symbols, respectively. In (F), the green arrow denotes position of male forelimb during grooming.

### GROSS SEXUAL BEHAVIORS LARGELY INTACT IN PEA3 NULL MALES

Next, a detailed analysis of gross sexual behaviors was performed to ascertain whether the inability of *Pea3* null mice to plug was due to a decreased propensity to mount receptive females. While there were fewer *Pea3* null males that mounted females, this difference was not statistically significant (Figure 2A). Of both control and *Pea3* null mice that did mount, we found that there was neither a difference in the time it took mice to perform their first mount (Figure 2B) nor in the number of mounts performed (Figure 2C). This indicates that the gross sexual motor behaviors are largely intact in *Pea3* null male mice.

**Figure 2.**
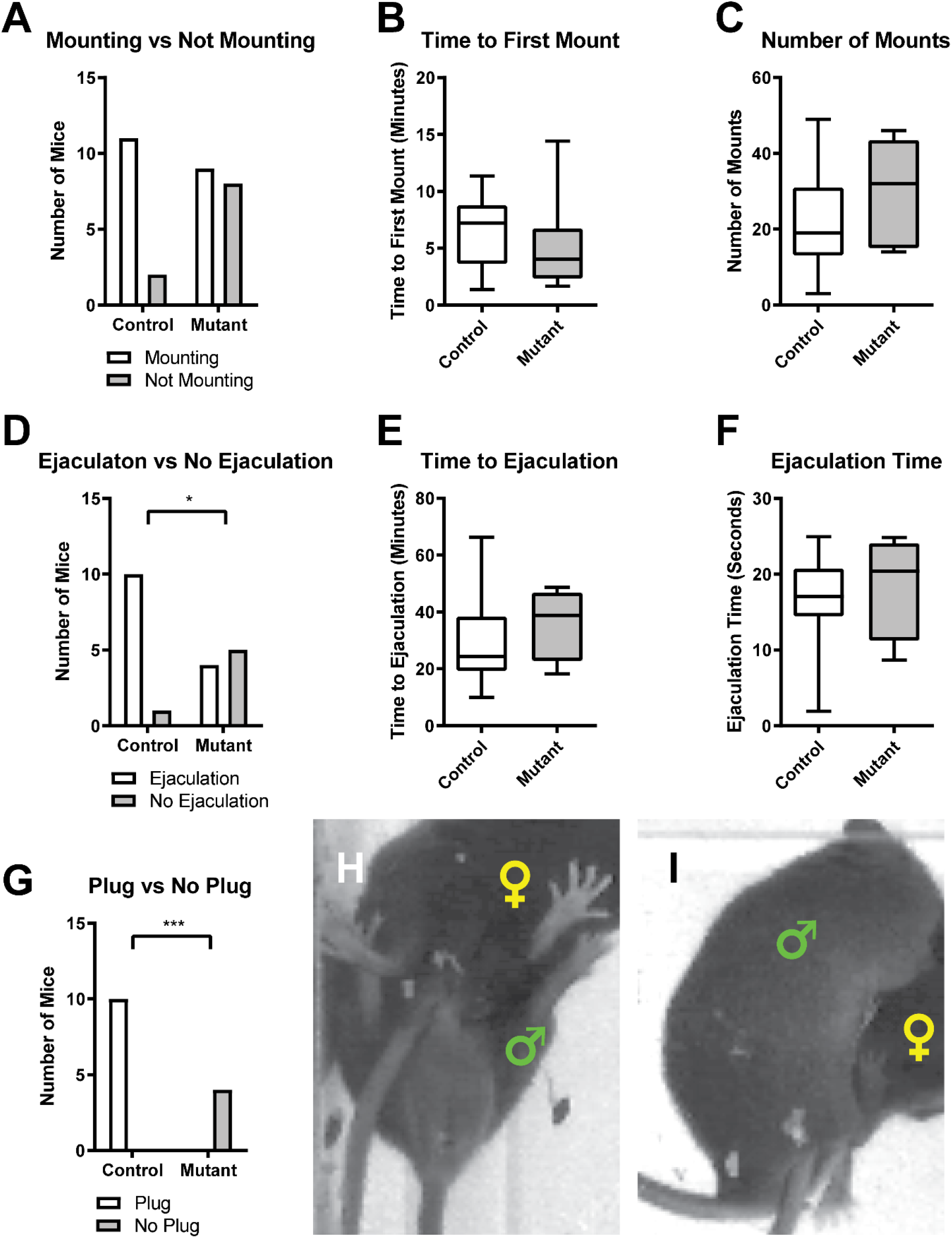
Quantification of gross sexual behavior deficits in *Pea3* null mice. (A) The number of mice that perform mounting behaviors is not statistically significant between control and *Pea3* null mice (control: N = 13 mice; null: N = 17 mice; Fisher’s exact test, p < 0.1194). (B and C) There is no difference between control and *Pea3* null mice in time to first mount (B; control: 6.32±0.93 minutes, n = 11; null: 5.05±1.37 minutes, n = 9; Student’s t-test, p < 0.4423) and the number of mounts performed (C; control: 21.45±4.25 mounts, n = 11; null: 29.44±4.49 mounts, n = 9; Student’s t-test, p < 0.2142). (D) Fewer *Pea3* null mice perform the ejaculatory grasping behavior compared to control mice (control: N = 11 mice; null: N = 9 mice; Fisher’s exact test, p < 0.0498, *). (E and F) There is no difference between control and *Pea3* null mice in time to ejaculatory grasping (E; control: 30.01±5.04 minutes, N = 10 mice; null: 36.16±6.47 minutes, N = 4 mice; Student’s t-test, p < 0.5084) and the duration of ejaculatory grasping (F; control: 16.41±1.97 seconds, N = 10 mice; null: 18.58±3.51 seconds, N = 4 mice; Student’s t-test, p < 0.5797). (G) Of mice that perform ejaculatory grasping behavior, no *Pea3* null mice deposit semen plugs compared to all control mice depositing semen plugs (control: N = 10 mice; null: N = 4 mice; Fisher’s exact test, p < 0.001, ***). Representative images of mounting (H) and ejaculatory behavior (I). Male and female positions are marked by green and yellow symbols, respectively.

One explanation for the loss of the ability to deposit semen plugs is that ejaculatory behavior was abolished *Pea3* null males^21,22^, therefore we assessed whether mice performed the stereotypical ejaculatory grasping behavior, during which the male stabilizes the female during insemination by grasping the female and falling over to one side. However, we found that *Pea3* null males were still capable of performing ejaculatory grasping behaviors, albeit at a much lower frequency than control males, thereby ruling out delayed or absent ejaculation-related behaviors (Figure 2D). Additionally, *Pea3* null mice that perform this ejaculatory grasping did so within the same amount of time and for the same duration as control males (Figure 2E and F), indicating the ejaculatory grasping behavior was of a sufficient time span necessary to deposit a semen plug.

Despite performing mounts and ejaculatory grasping behavior, we confirmed that *Pea3* null males fail to deposit semen plugs within the vagina (Figure 1G), as previously reported^16^. Additionally, we did not witness expulsion of semen from the penis of *Pea3* null mice at any time during our mating assays, therefore, the lack of plug deposition was not due to aberrant extra-vaginal placement of expelled semen.

### *PEA3* NULL MALE MICE EXHIBIT GREATLY DIMINISHED ERECTILE FUNCTION DURING SEXUAL INTERCOURSE

Since *Pea3* null males display sufficiently normal gross mating behavior, yet fail to inseminate females, we next focused our analysis on the placement of the penis and the ability to achieve erection. The inability to deposit semen plugs may derive from two possible sources: (1) perturbations in ejaculatory abilities that leave overall erectile functioning intact^21,22^, or (2) erectile dysfunction that decreases the ability to achieve or maintain an erection. Monitoring erectile function in the copulating mouse is challenging because of the speed with which males transition from the initiation of the mount to intromission, however, by replaying recordings of mounting bouts at ¼ speed, we found that erectile function in *Pea3* null male mice is dramatically reduced. *Pea3* null males do not produce visible erections during the vast majority of mounting bouts (Figure 3A-C, G; supplemental video 1). During the rare mounting bouts when the penis exits the penile sheath, it is only briefly (∼0.1 seconds) visible at end of trains of high frequency thrusts and is not properly targeted to the vagina to allow for intromission. In contrast, control males generate and maintain visible erections during the initiation of mounts and high frequency thrusting that immediately precedes intromission (Figure 3A-C and F; supplemental video 2). As such, *Pea3* null mice fail to achieve intromission, both when quantified as the overall number of intromissions and intromissions per mount (Figure 3B and C). Therefore, the erectile deficiencies present in *Pea3* null males offer the clearest explanation for their inability to deposit semen plugs in receptive females.

**Figure 3.**
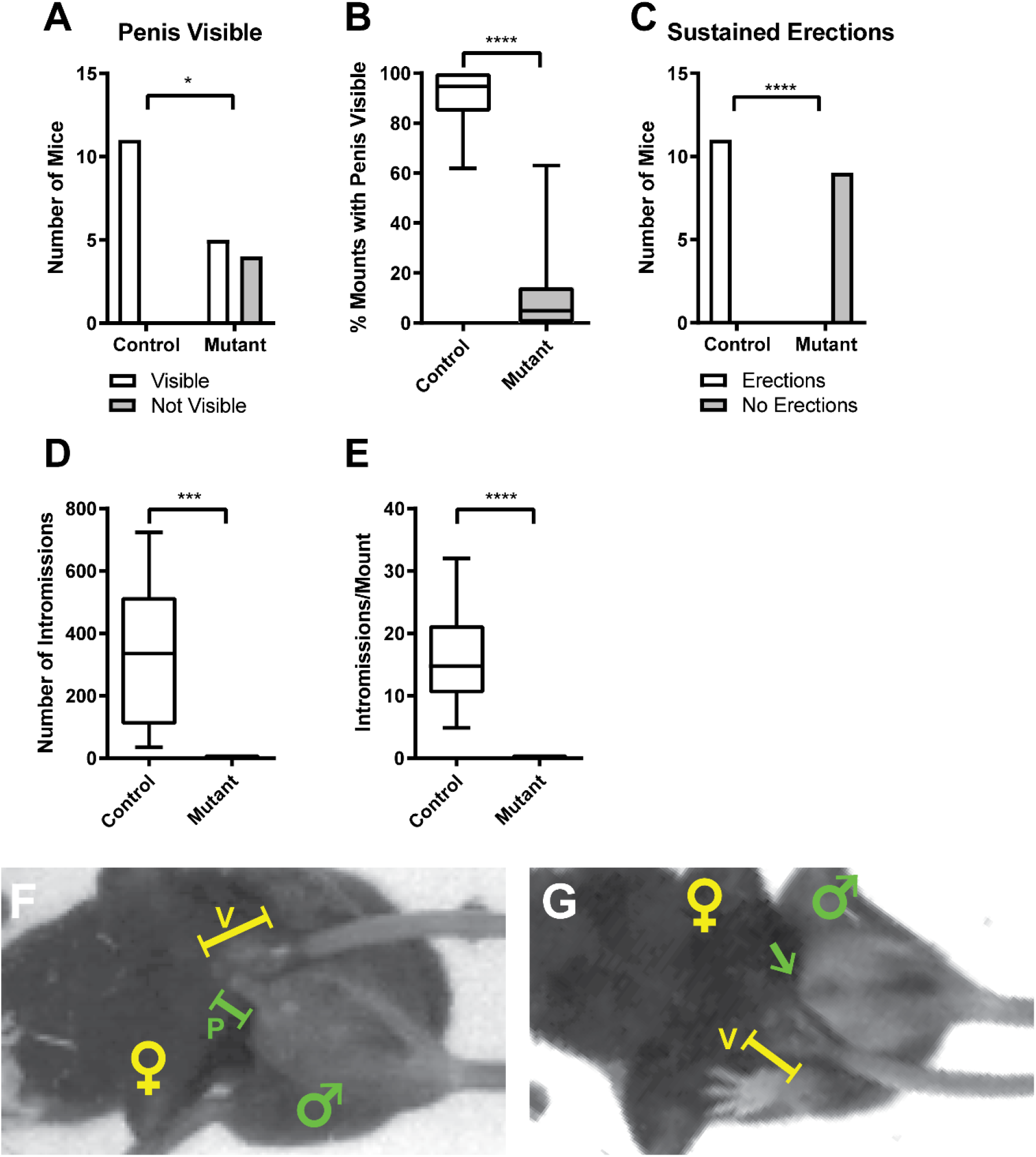
*Pea3* null males have greatly diminished erectile functioning during sexual behavior. (A) Of mounting mice, there are fewer *Pea3* null mice with the penis visible outside of the penile sheath compared to control mice (control: N = 11 mice; null: N = 9 mice; Fisher’s exact test, p < 0.026, *). (B) *Pea3* null mice exhibit a decreased percentage of mounts with the penis visible outside of the penile sheath during mounting behavior compared to control mice (control: 89.60±3.871%, N = 11 mice; null: 11.83±6.715%, N = 9 mice; Student’s t-test, p < 0.0001, ****). (C) Of mounting mice, no *Pea3* null mice exhibit sustained erections during sex as opposed to all control mice doing so (control: N = 11 mice; null: N = 9 mice; Fisher’s exact test, p < 0.0001, ****). (D and E) *Pea3* null mice do not achieve intromission, as compared to control mice for the total number of intromissions performed (D; control: 331.20±68.32 intromissions, N = 11 mice; null: 0.0±0.0 intromissions, N = 9 mice; Student’s t-test, p < 0.0004, ***) and the number of intromissions per mount (E; control: 16.71±2.75 intromissions/mount, N = 11 mice; null: 0.0±0.0 intromissions/mount, N = 9 mice; Student’s t-test, p < 0.0001, ****). Representative images of the presence (control mice; F) or absence (*Pea3* null mice; G) of erectile function. Male and female positions are marked by green and yellow symbols, respectively. Green capped lines and arrows indicate the position of the penis (P), and yellow capped lines indicate the position of the vagina (V).

### PERTURBED INTROMISSION-LIKE THRUST TIMING IN PEA3 NULL MALE MICE

The abolishment of erectile functioning in *Pea3* null mice, despite otherwise largely intact gross sexual behaviors, offers an opportunity to address how sensory feedback from intravaginal penile contact influences the number, timing, and variability of hip thrusts during intromission. By definition, intromission implies the insertion of the penis into the vagina, therefore for *Pea3* null mice we use the term “intromission-like” to describe various components of intromission associated behaviors (i.e., count, timing, etc.). Taking advantage of the high temporal resolution of our data recordings, we classified intromissions and intromission-like behaviors by the inter-thrust timing, or the time it takes from the start of one hip thrust to the start of the next hip thrust. These data were collected for both the first and last mounting sequence recorded for each mouse during the sexual behavior assay. For both the first (Figure 4A and B) and last (Figure 4C and D) mounting sequence, *Pea3* null mice have a clear shift in intromission-like thrust timing toward shorter duration intromission thrust times compared to control mice. Interestingly, the number of intromission-like thrusts was not different between control and *Pea3* null males, for either the first or last mounting sequence (Figure 4E and F). These data indicate that intravaginal contact is not necessary to initiate a train of intromission-like thrusts.

**Figure 4.**
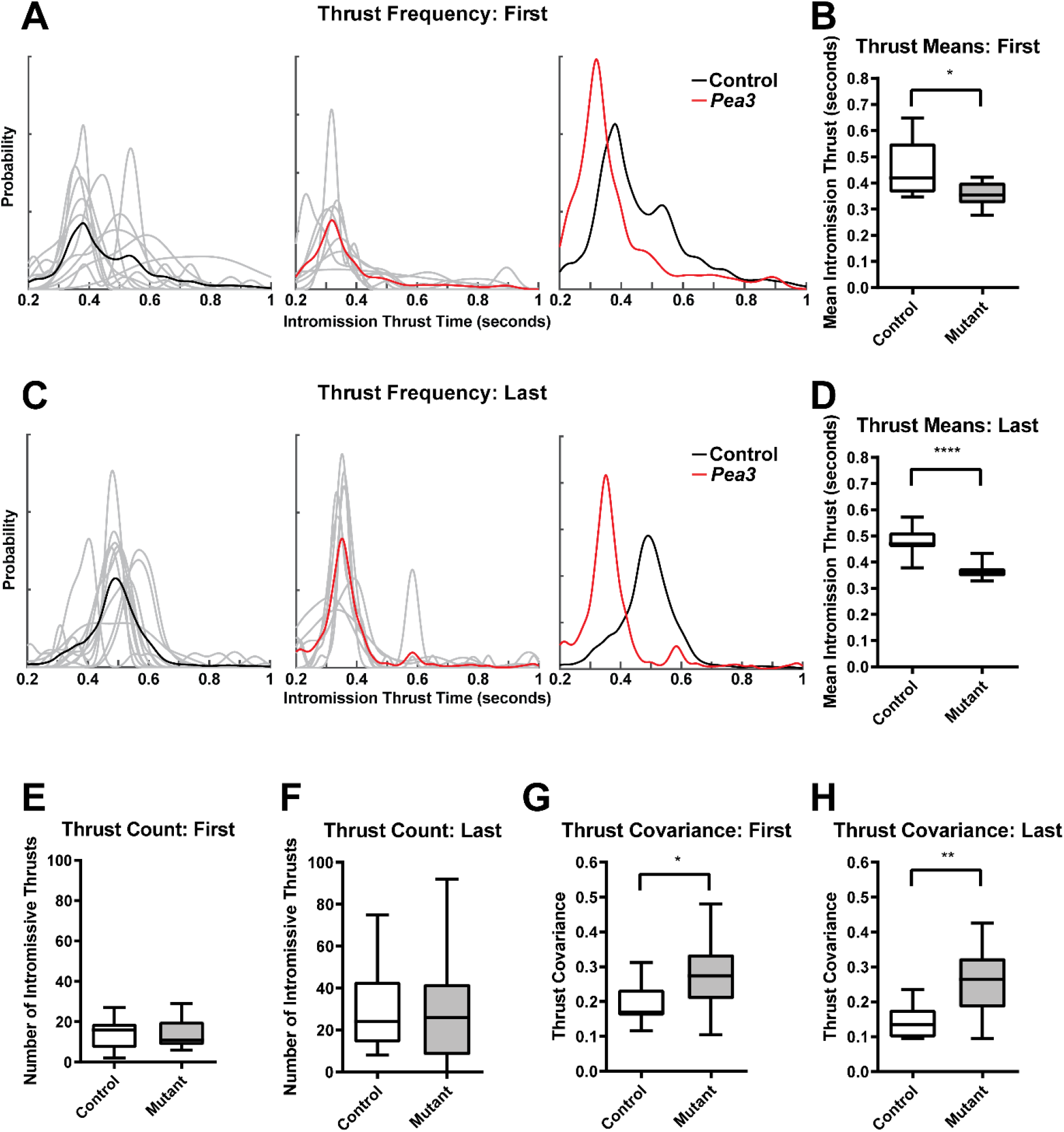
Quantification of shifts in intromission-like thrusting behavior in *Pea3* null males. (A and C) The timing of intromission-like thrusting shifts for both the first (A) and last (C) mounts between control (black line; left and right panels) and *Pea3* null (red line; middle and right panels) males. Solid colored (black and red) lines are produced by averaging the distributions from individual animals (gray lines; left and middle panels). (B and D) Intromission-like thrusts are faster in *Pea3* null as compared to control males for both the first (B; control: 0.452±0.030 seconds, N = 11 mice; null: 0.359±0.016 seconds, N = 9 mice; Student’s t-test, p < 0.0191, *) and last (D; 0.483±0.017 seconds, N = 11 mice; null: 0.366±0.011 seconds, N = 8 mice; Student’s t-test, p < 0.0001, ****) mount. (E and F) The number of intromission-like thrusts does not change between control and *Pea3* null males for the first (E; control: 14.27±2.37 thrusts, N = 11 mice; null: 14.11±2.49 thrusts, N = 9 mice; Student’s t-test, p < 0.9632) and last (F; control: 29.27±6.37 thrusts, N = 11 mice; null: 29.11±9.40 thrusts, N = 9 mice; Student’s t-test, p < 0.9885) mounts. (G and H) The coefficient of variation for thrust timing increases for *Pea3* null vs control males for both the first (G; control: 0.190±0.017, N = 11 mice; null: 0.278±0.036, N = 9 mice; Student’s t-test, p < 0.031, *) and last (H; control: 0.145±0.015, N = 11 mice; null: 0.258±0.036, N = 8 mice; Student’s t-test, p < 0.0051, **) mount.

Lastly, we observed that the thrusting trains in *Pea3* null males were highly disordered when compared to control males. In *Pea3* null males, fast thrusting patterns (less than 200 milliseconds) were interspersed between intromission-like thrusting patterns, whereas control males transitioned into stable intromission-like thrusting patterns after an initial bout of fast thrusting. We measured the variability of timing between adjacent hip thrusts in the thrusting sequence for both the first and last mounts by calculating the coefficient of variation (CoV) for thrust timing. Mice with more variability in their thrust times will have higher CoV values, and mice with highly similar thrust times will have lower values. For both the first and last mount, we found that there was a higher variability in thrust timings in *Pea3* null male mice versus control mice, indicating that *Pea3* null mice are less able to generate consistently timed intromission-like thrusts (Figure 4G and H).

## DISCUSSION

Here, we set out to assess *Pea3* null male mice for an array of potential deficiencies in male sexual behaviors that may occur during active copulation. We employed a sexual behavior arena that allows for the viewing of mounts, thrusts and intromissions from the mouse underside, granting a clear view of the penis during sexual activity^18^. We coupled this behavioral assay with high-speed, high-resolution video recording that allows for a detailed post-hoc analysis of sexual behaviors. Surprisingly, we found that *Pea3* null male mice have greatly diminished erectile function during copulation, and therefore never achieve intromission, despite displaying sufficiently normal gross motor behaviors associated with sexual intercourse (i.e., mounting, thrusting, and ejaculatory behavior).

There are few comparable studies within the literature that describe genetic mutations with similarly severely diminished erectile functioning, highlighting the novelty of the *Pea3* null male phenotype. The closest phenotypic match to *Pea3* null males are *TGF1beta* null male mice, which are infertile and do not deposit semen plugs during sexual behavior assays^10^. However, *TGF1beta* null males are able to achieve erections naturally (via self-grooming) despite clear deficits in the structural integrity of the penis^11^, an issue which is not present in *Pea3* null males^16^. Additionally, there are many other genetic mutations in mice that lessen or abolish the deposition of semen plugs during short, monitored copulation assays, however, do not render males infertile in paired mating assays of longer duration (weeks to months)^6,8,9,12,15,23–25^, indicating the presence of sufficient erectile function to reproduce, again, in contrast to *Pea3* null males^16^. The transcription factor *SIM1* has recently been linked to erectile dysfunction in humans^4^, but in mouse models the direct knockout of the *SIM1* gene has yet to be assessed for erectile dysfunction. Homozygous *SIM1* knockout mice die perinatally^26^ and postnatal deletion of *SIM1* in mice leads to hyperphagic obesity^27^, complicating the search for an erectile deficiency directly linked to altered *SIM1* expression. This highlights the novelty of the *Pea3* mutant phenotype, as homozygous *Pea3* null male mice are viable, display largely unaltered appetitive and consummatory male sexual behaviors, yet display greatly diminished erectile function during sexual intercourse. Interestingly, *Pea3*, also known as *Etv4*, has been linked to human urological cancers, however, has not been directly investigated as a cause for erectile dysfunction^28^.

How could *Pea3* affect erectile function? As *Pea3* is expressed within the brain and spinal cord^16,17^, our data suggest a possible role for *Pea3* in organizing motor circuits that control the pudendal muscles, the bulbocavernosus (BC) and ischiocavernosus (IC). For wildtype males, the BC and IC muscles contract synchronously during intromission, which increases intracavernosal pressure to achieve the erectile rigidity necessary for vaginal insertion^29^. During ejaculation, the activity of the BC and IC muscles shifts from synchronous to alternating, which turns the urethra into a pump and drives the expulsion of semen^29^. After excision of the IC muscle, successful intromissions are greatly reduced, however males continue to mount, thrust, and display ejaculatory patterns despite their markedly reduced ability to achieve intromission^30^. Excision of the BC muscle has little effect on erectile function, but reduces fertility during mating assays, which is suggested to be due to improper plug placement during ejaculation^31^. Similarly, after transection of the motor branch of the pudendal nerve, which supplies motor efferent innervation for IC and BC muscles, there is a decrease in intromissions and ejaculations, a large increase of extravaginal intromission patterns, and an inability to gain erections during reflexive tests^32^. The phenotype of *Pea3* null male mice is similar to that produced by the excision of the IC and BC muscles, or the transection of motor afferents that supply them, namely, greatly diminished erectile function and extravaginal penile targeting during thrusting behaviors. Secondarily, the brief and poorly timed periods of penile tumescence witnessed in *Pea3* null males suggest insufficient control over the timing of increases in intracavernosal pressure necessary for intromission, which is governed by the pudendal muscles and upstream sexual motor circuitry in the spinal cord.

The outcomes of our study were dependent on our choice of a sexual behavior arena with a clear floor and an angled mirror placed underneath, which allows for the unobstructed viewing of the underside of mice during sexual behavior. This apparatus, elegantly detailed in Park (2011)^18^, has been employed for many decades^33^. The privileged viewing angle possible with this arena, when coupled with digital videography (as described in this study), allows for unambiguous identification of erectile functioning, including penile tumescence and the spatial relationship of the penis relative to the vagina. The clear view of the penis during sexual behaviors abrogates the necessity of inferring erectile function either using reflexive assays^8,11,15,16^, or by monitoring hip thrusting patterns of the copulating male mouse^7,8,10–13,25,34^. Increased adoption of this behavioral arena when monitoring male sexual behaviors will aid in further identifying erectile deficiencies in previously characterized and novel mouse models of male sexual dysfunction. While outside of the goals for this current study, this setup is well suited for combining sexual behavioral analysis with deep learning algorithms. Such an approach will facilitate screening mutant mouse lines to identify novel genes associated with erectile function and greatly advance the study of sexual behaviors and erectile functioning^35–37^.

Lastly, many studies have sought to identify the role of sensory feedback due to successful intromissions on the organization and execution of mouse sexual motor behaviors, either by surgically inducing impotence^30,32,38^, the application of potent topical local anesthetics to the penis^38–41^, or by limiting access to the vagina^39,42^. The erectile deficiencies witnessed in *Pea3* null male mice offers an opportunity to address the role of erectile sensory feedback genetically, without need for pre-testing interventions, surgical or otherwise. Our data show that there are alterations in hip thrusting frequency patterns in *Pea3* null males, potentially due to lost sensory feedback from intravaginal insertion during intromission. As such, the *Pea3* null mouse line is a valuable genetic tool with erectile deficits beyond the previously identified deficits in fertility and ejaculatory function.

Future studies are necessary to interrogate the molecular mechanisms of *Pea3* in controlling erectile function. Such insights could potentially aid in the discovery of treatments for erectile dysfunction in patients^3^.

## ACKNOWLEDGMENTS

We are grateful to Silvia Arber (FMI Basel) for suggesting that we assay *Pea3* mutants for sexual behavior phenotypes, Nirao Shah (Stanford University) for providing technical advice and comments on this manuscript, Jin Ho Park (University of Massachusetts, Boston) and Dayu Lin (New York University) for providing technical advice on setting up sexual behavior assays, Bill Carson (Systematic Vision, Inc.) for advice on digital recording apparatus design, and the carpentry staff at MSKCC and Jarret AT Weinrich for aiding in the construction of behavioral arenas. This work was supported by Memorial Sloan-Kettering start-up funds, MSK Cancer Center Support Grant/Core Grant (P30 CA008748), a Louis V. Gerstner, Jr. Young Investigators Award, and NIH grant R01 NS083998 (J.A.K.).

