## supplemental for "Transcription Factor Gene *Pea3* Regulates Erectile Function in Mice"

**Supplemental Video 1: *Pea3* null males have greatly diminished erectile functioning during sexual behavior.**

*Pea3* null males fail to generate erections during active copulation. Mounting and thrusting behavior is displayed at 1X and 1/10X video playback speed.

Link to Supplemental Video 1: <https://ucsf.box.com/v/2021-Pea3-Supp-Vid-01>

**Supplemental Video 2: Control males exhibit normal erectile functioning**

Control males generate erections during active copulation. Mounting and thrusting behavior is displayed at 1X and 1/10X video playback speed.

Link to Supplemental Video 2: <https://ucsf.box.com/v/2021-Pea3-Supp-Vid-02>
